# Voluntary adolescent alcohol exposure does not robustly increase adulthood consumption of alcohol in multiple mouse and rat models

**DOI:** 10.1101/2024.04.30.591674

**Authors:** Avery R. Sicher, Andrea Liss, Valentina Vozella, Paige Marsland, Laurel R. Seemiller, Matthew Springer, William D. Starnes, Keith R. Griffith, Grace C. Smith, Amy Astefanous, Terrence Deak, Marisa Roberto, Florence P. Varodayan, Nicole A. Crowley

## Abstract

Adolescence is a period of increased risk taking, including increased alcohol and drug use. Multiple clinical studies report a positive relationship between adolescent alcohol consumption and risk of developing an alcohol use disorder (AUD) in adulthood. However, few preclinical studies have attempted to tease apart the biological contributions of adolescent alcohol exposure, independent of other social, environmental, and stress factors, and studies that have been conducted show mixed results. Here we use several adolescent voluntary consumption of alcohol models, conducted across four labs in three institutes and with two rodent species, to investigate the ramifications of adolescent alcohol consumption on adulthood alcohol consumption in controlled, pre-clinical environments. We consistently demonstrate a lack of robust increases in adulthood alcohol consumption. This work highlights that risks seen in both human datasets and other murine drinking models may be due to unique social and environmental factors – some of which may be unique to humans.

**HIGHLIGHTS:**

- Adolescent drinking-in-the-dark (DID) binge drinking does not increase adulthood consumption in a DID model or a two bottle choice model in male and female SST-Cre:Ai9 mice
- Adolescent pair-housed intermittent access consumption of alcohol does not increase adulthood consumption in an identical adulthood model in male and female C57BL/6J mice
- Adolescent intermittent access to alcohol does not increase adulthood consumption in male and female Wistar or Fischer 344 rats
- These complementary datasets across murine models, labs and institutions highlight the need to consider human social factors as well as biological factors

## 1.0 INTRODUCTION

Adolescence marks the transitional period between childhood and adulthood (Spear, 2000), with initiation of substance use often beginning during this time. This increase in substance use including alcohol poses both immediate risks (car accidents, violence, unsafe intercourse) and long-term risks (chronic organ damage, neuropsychiatric conditions, and alcohol use disorders) (Chung et al., 2018; Vaca et al., 2020). Several studies have reported a relationship between age at first alcohol use and an increased risk of developing an alcohol use disorder (AUD) later in life (Grant & Dawson 1997; DeWit et al., 2000; Addolorato et al., 2018), although other analyses suggest age at first intoxication or the onset of regular drinking episodes may be a more reliable predictor of subsequent risky drinking patterns (Sartor et al., 2016; Newton-Howes, et al., 2019). However, a variety of additional environmental factors and general risk factors likely influence both the age of first alcohol use onset and AUD risk. There is also an increasing need for in-depth investigation into sex- and gender-based differences in alcohol consumption, as rates of binge drinking in adolescent girls and boys continue to converge, driven by slower declines in binge drinking rates in adolescent girls (White, 2020).

Understanding the risk factors for escalated drinking could lead to the development of improved and more individualized treatment strategies for problematic alcohol use in adults. Animal models can be used to further tease apart biological and environmental contributions in a causative manner; however, the relationship between early life alcohol exposure and alcohol consumption in adulthood is often inconsistent in these studies. A positive relationship between adolescent alcohol exposure and subsequent adult alcohol consumption has been reported in several rodent studies using a variety of exposure paradigms (Strong et al., 2010; Younis et al., 2019; Wolstenholme et al., 2020). However, this effect is not universally observed (Vetter et al., 2002; Gilpin et al., 2012; Wooden et al., 2023). Factors which vary between studies and may contribute to the contrasting findings include sex and strain of the rodents used, the precise age window of early alcohol exposure, models of alcohol exposure (i.e., voluntary versus forced), and housing conditions (for review, see Towner and Varlinskaya, 2020).

In this series of experiments, we used four different paradigms of voluntary adolescent alcohol exposure across two species (mouse and rat) and three institutional sites (Penn State University, Binghamton University - SUNY, and The Scripps Research Institute) and examined changes in alcohol consumption and preference in adulthood using multiple models. The first two experiments used the Drinking-in-the-Dark (DID) paradigm in mice, modified for adolescent timepoints, coupled with either 4 additional weeks of DID during adulthood or 6 weeks of a continuous access 2-bottle choice procedure (2BC). The third experiment used a model of intermittent access 2-bottle choice (IA2BC) beginning in adolescence with an identical model during adulthood in mice. The fourth experiment used an intermittent access design in rats. The final experiment used a model of chronic intermittent ethanol (CIE) drinking in adolescence and adulthood in rats. All experiments began alcohol exposure in early adolescence, typically thought to begin in the fourth postnatal week (∼ PND 28) of rats and mice, while late adolescence spans into the eighth postnatal week (∼ PND 56; Spear, 2015; Schneider, 2013; Bell, 2018). All of these models are well validated for their ability to yield high levels of alcohol consumption during adolescence (Sicher et al., 2023; Melendez, 2011, Marsland et al., 2023A), and complement each other across model-induced stressors, individual data, and drinking patterns. Based on the previous literature, we hypothesized that adulthood alcohol consumption would be increased across all four models of early-adolescent voluntary drinking. However, we do not see consistent changes in adult drinking following these complementary models of adolescent binge and intermittent drinking.

## 2.0 MATERIALS AND METHODS

### 2.1 Animals

Experiments were approved by the Pennsylvania State University Institutional Animal Care and Use Committee (University Park, PA), the Binghamton University Institutional Animal Care and Use Committee (Binghamton, NY), and The Scripps Research Institute Animal Care and Use Committee (San Diego, CA). All experiments used voluntary consumption models beginning in early adolescence (for review, see Crowley et al., 2019).

For experiments conducted at Penn State (**Figures 1, 2, 3**), 77 male and female SST-IRES-Cre:Ai9 mice on a C57BL/6J background were bred in-house from SST-IRES-Cre (stock #0130444, The Jackson Laboratory) and Ai9 (stock #007909, The Jackson Laboratory) mice. SST-IRES-Cre:Ai9 mice express a fluorescent reporter only in somatostatin-expressing neurons, which was used for other experiments not reported in this paper and only after drinking was completed. Upon weaning at postnatal day (PND) 21, mice were single housed and moved into a temperature- and humidity-controlled reverse light cycle room (lights off at 7:00 am). Mice acclimated for one week prior to the start of adolescent drinking experiments. Mice had *ad libitum* access to food and water, except during limited access alcohol exposure outlined below. Throughout all experiments, mice were weighed once weekly.

For Experiment 4 conducted at Binghamton University (**Figures 1, 4**) 62 male and female C57BL/6J mice were bred in-house in a temperature- and humidity-controlled reverse light cycle room (lights off at 10:00 am). Following weaning at PND 21, mice were pair-housed with same-sex littermates and then left undisturbed until the start of adolescent drinking (PND 30). All mice had *ad libitum* access to food and water. All mice were weighed once a week throughout the duration of all drinking experiments. Only one pair of mice per sex per litter were included in each experimental group.

For Experiment 5, conducted at The Scripps Research Institute (**Figures 1, 5**), 14 male and female Wistar rats were purchased from Charles River Laboratories (Wilmington, MA, USA) and shipped to Scripps’ animal facility upon weaning at postnatal day PND 21. On their arrival, rats were group-housed (4/cage) and moved into a temperature- and humidity-controlled reverse light cycle room (lights off at 8:00am). Rats acclimated for 5 days prior to the start of adolescent drinking experiments. Rats had *ad libitum* access to food and water. Throughout the experiment, rats were weighed after each drinking session.

For Experiment 6, conducted at Binghamton University (**Figures 1, 6**), Fischer 344 rats were acquired from Charles River and bred on site. At PND 21, rats were weaned and pair-housed with a same-sex, non-littermate partner for experimentation prior to PND 28. Experimental subjects had *ad libitum* access to food and were housed in standard Plexiglass cages, with wooden chew blocks provided as enrichment. Colony conditions were maintained at 22 +/- 1° C with a 12-hr light-dark cycle (lights on 7:00am).

### 2.2 Adolescent Drinking in the Dark (DID) in mice at Penn State

Adolescent DID was conducted as previously published (Sicher et al., 2023; **Figure 1A-D**). Starting between PND 28-30, mice received 20% (w/v) ethanol (EtOH; Koptec, Decon Labs, King of Prussia, PA) diluted in tap water in a sipper tube beginning 3 hours into the dark cycle for 2 hours on 3 consecutive days (i.e. 10:00am to 12:00pm; Rhodes et al., 2005). On the fourth day, mice had access to EtOH for 4 hours (i.e. 10:00am to 2:00pm), representing the “binge” day. Sipper tubes were weighed at the start and end of each drinking period and EtOH intake was normalized to body weight. After the binge day, mice had 3 days of abstinence without any ethanol access. This cycle was repeated for a total of 4 weeks. During adolescent DID, control mice had access to water only.

Blood samples via cardiac puncture were collected within 30 minutes of the final binge session to measure blood ethanol concentration (BEC) in a subset of mice. BECs were determined using an Analox AM1 analyzer to confirm binge-like consumption (Analox Instruments, Lunenberg, MA). For all other mice, following the end of adolescent DID (PND 52-54), mice sat undisturbed except for weekly cage changes and weighing for approximately 30 days. At PND 84, mice were assigned to either 4 weeks of adulthood DID (experiment 1) or 6 weeks of adulthood 2-bottle choice (experiment 2), as described below.

#### Experiment 1: Adulthood Drinking in the Dark (“double DID”)

Beginning at PND 84, all mice assigned to the adulthood DID experiment began 4 weeks of DID as previously described (**Figure 2**). In this experiment, all mice (e.g., both those that had undergone adolescent DID and control conditions) had access to 20% EtOH via the DID paradigm for four weeks.

#### Experiment 2: Continuous Access Adulthood 2-Bottle Choice (2BC)

A subset of adolescent DID and control exposed mice were used for adulthood 2-bottle choice (2BC; **Figure 3**). One week prior to the start of 2BC, at PND 77, mice were given continuous access to one sipper tube filled with normal tap water for acclimation and tracking of baseline fluid consumption. After acclimation, all mice underwent 6 weeks of 2BC as described previously (Dao et al., 2020). Mice received continuous access to one sipper tube of tap water and one sipper tube of unsweetened ethanol diluted in tap water. Ethanol concentrations were increased from 3% (w/v) for days 1-3, 7% for days 3-9, and 10% for days 9-42. During 2BC, bottles were weighed and refilled every 48 hours, with the positions of the bottles alternated each cycle. Preference was calculated as the amount of ethanol solution consumed per 48-hour period divided by the total fluid consumption over the same time.

### 2.3 Pair-housed intermittent access 2-bottle choice (IA2BC) in mice at Binghamton University

Starting on PND 30, adolescent pair-housed littermates underwent either intermittent access two bottle choice (IA2BC) drinking or water drinking only (**Figure 1E-G**). During the first day (PND 30), pair-housed cages that underwent IA2BC had access to one bottle of water and one bottle of 3% (v/v) unsweetened EtOH diluted in tap water for 24 hours. For the next 24 hours, mice had access to two bottles of tap water. This cycle repeated with mice having access to increasing concentrations of EtOH (2 days of 3%, 1 day of 6%, 1 day of 10%), building to a final concentration of 15% ethanol used for remaining time. A total of 15 EtOH drinking sessions were completed, and adolescent drinking ended at PND 60. Control mice had access to two bottles of water from PND 30-60. Since mice were pair-housed throughout their EtOH drinking sessions, daily EtOH and water intake were measured per cage and divided by the combined body weight of the pair-housed mice. As a result, each individual data point represents a cage of two same-sex litter-mate mice. Similarly, daily EtOH preference was calculated per cage as the volume of ethanol consumed divided by the total fluid consumed during each drinking session.

#### Experiment 3: Adulthood intermittent access 2-bottle choice (IA2BC)

For adult drinking, all pair-housed cages underwent IA2BC regardless of whether they underwent EtOH drinking during adolescence or water drinking only (**Figure 4**). A similar IA2BC as described above was employed, however the mice only had access to 15% EtOH throughout the duration of adult drinking. Mice drank from PND 90-120 for a total of 15 EtOH drinking sessions.

### 2.4 Intermittent access 2-bottle choice (IA2BC) in rats at The Scripps Research Institute

At PND 28-30, male and female Wistar rats started receiving access to water and 20% (v/v) EtOH (Pharmco by Greenfield Global, Shelbyville, KY) in tap water in a sipper tube beginning 2 hours into the dark cycle for 2 hours every other day (i.e. 10:00am to 12:00pm; Tuesday/Thursday/Saturday; **Figure 1H-K**). 15 minutes prior to IA2BC, rats were moved to a mating cage divided with perforated clear plastic dividers to ensure each animal had access to its own water and EtOH bottle while reducing isolation stress. Sipper tubes were weighed at the start and end of each drinking period. The location of drinking tubes was switched at each drinking session to avoid development of side preference. This cycle was repeated for 4 weeks.

#### Experiment 4: Adulthood intermittent access 2-bottle choice (IA2BC)

During the 2 weeks of alcohol abstinence, rats were left undisturbed, housed in 4/cage and given continuous access to two sipper tubes filled with normal tap water. At PND 70, rats underwent 6 sessions of 2 hours IA2BC, every other day, as in adolescence (**Figure 5**).

### 2.5 Chronic intermittent ethanol exposure (CIE) in rats at Binghamton University

Adolescent (PND 28-PND 32) Fischer 344 rats were placed on an intermittent EtOH exposure procedure (**Figure 1L-N**). Rats were given access to a single bottle of EtOH (10% v/v in tap water) for a period of 48 hours, followed by a period of only tap water access for 48 hours (two days on/two days off). Control rats had access to a single bottle of tap water only. Each presentation of EtOH followed by water was termed a cycle, and rats were placed on this procedure for a total of 12 cycles (48 days; ending at PND 74) during adolescence. Bottles of either EtOH or water were weighed daily, and g/kg intake was estimated within the cage using the combined bodyweights of the paired rats. Each data point represents one cage of 2 rats.

#### Experiment 5: Chronic intermittent ethanol exposure (CIE)

Following a period of abstinence (PND 74-114), adult animals were then placed back on the CIE exposure procedure for another 12 cycles (**Figure 6**). Data from our lab using this model indicates that the procedure is well tolerated and produces consistent, moderate levels of EtOH consumption based on both blood and brain EtOH concentrations (Marsland et al., 2023A; Marsland et al., 2023B).

### 2.5 Statistics

Statistics were run in GraphPad Prism and Matlab. Two and three-way ANOVAs (adolescent EtOH condition, sex, and drinking session, where applicable) were used for between-group comparisons, while generalized linear modeling was used to explore data more deeply in adolescent-EtOH mice only. For ANOVAs, data were matched within each subject when appropriate to account for repeated drinking sessions (e.g. within-subject comparisons, comparisons of multiple binge days). Effect sizes for ANOVAs were calculated in Prism and reported as ω^2^.

Across all alcohol-only datasets, a generalized linear mixed-effects model (GLME) accounting for litter as a random effect variable was used to analyze drinking in adulthood. We included litter as a random effect in the GLME for experiments 1 and 2, as multiple subjects were taken from each litter. The model was as follows: [outcome variable] ∼ 1 + Sex * Total adolescent EtOH + (1|Litter). For Experiment 1 and 5, measuring adulthood EtOH intake only, the GLM used sex and total adolescent EtOH intake as predictors for each outcome variable. Outcome variables included EtOH intake during the first and last adulthood drinking sessions and total EtOH intake during each adulthood paradigm. For all experiments using 2BC (Experiments 2, 3, and 4), outcome variables included EtOH preference and EtOH intake during the first and last adulthood drinking sessions and total EtOH consumed across drinking sessions. For preference during a 2BC paradigm (Experiments 2, 3, and 4), the GLM for preference used the Logit link function. We chose these outcome variables to give a broad understanding of changes in drinking patterns following adolescent EtOH.

Outliers in daily drinking (due to infrequent bottle leaks; experiments 1 and 2) were determined as a bottle change of more than 2g in 2 hours, matching outliers identified the ROUT outlier test in GraphPad Prism (Q = 1%). These days were removed from daily drinking analyses. For total EtOH consumption, a bottle leak was replaced by the average consumption of other days that week (so that equal numbers of drinking days were included in the total consumption).

## 3.0 RESULTS

### 3.1 Overall adolescent binge drinking models and consumption levels demonstrate all models lead to robust alcohol consumption

In experiments 1 and 2, male and female SST-Cre:Ai9 mice underwent 4 cycles of DID or water control beginning between PND 28-30 (**Figure 1A-D**). In the mice which underwent adolescent EtOH DID, a repeated-measures ANOVA (factors: sex, binge day) showed a main effect of sex on binge alcohol consumption (across 4h binge days; **Figure 1B**; *F*_sex_(1, 36) = 10.84, *p* = .0022; *F*_binge day_(2.393, 82.95) = 0.2687, *p* = .8031; *F*_sex x binge day_(3, 104) =1.205, *p* = .3119). Post-hoc testing using Šídák’s multiple comparisons test revealed that female mice consumed more EtOH than male mice during the second adolescent binge day (PND 38; *t*(29.01) = 3.254; adjusted *p* = 0.0115; for all other adolescent binge days, *p* > .05). An unpaired t-test showed a sex difference in total EtOH consumption across all 16 days of adolescent DID (**Figure 1C**; *t*(36) = 3.027; *p* = .0045). Across all 16 days of adolescent DID, female mice drank more EtOH than male mice. An F-test to compare the variances in total ethanol adolescent alcohol consumption showed no differences in the variances between male and female mice (*F*(20, 16) = 1.634; *p* = .3227). BECs were assessed immediately following the final adolescent binge day (PND 52) to confirm binge-levels of consumption (**Figure 1D**).

In experiment 3, male and female C57BL/6J mice underwent intermittent access 2-bottle choice (IA2BC) protocol beginning at PND 30 (**Figure 1E-G**). Within mice that underwent EtOH IA2BC during adolescence, a 2-way ANOVA (factors: sex and drinking session) indicated an interaction between sex and drinking session (**Figure 1F**; *F*_sex x session_(14, 182) = 3.869; *p* < .0001). Šídák’s multiple comparisons test did not indicate significant differences in EtOH intake in male and female mice during any individual drinking session (*p* > .05). Similarly, there was a sex x drinking session interaction in preference for EtOH during adolescent IA2BC (**Figure 1G**; *F*_sex x session_(14, 182) = 1.976; *p =* .0218). Šídák’s multiple comparisons test did not indicate significant differences in EtOH preference in male and female mice during any individual drinking session (*p* > .05).

In experiment 4, male and female Wistar rats underwent an IA2BC paradigm starting at PND 28 (**Figure 1H-K**). A 2-way ANOVA (factors: sex and drinking session) revealed a significant main effect of drinking session on 2-hour ethanol intake (**Figure 1I**; *F*_drinking session_(3.378, 40.54) = 2.757; *p* = .0487), but not of sex (*F*_sex_(1, 12) = 0.008907; *p* = .9264) nor an interaction (*F*_sex x drinking session_(11, 132) = 1.194; *p* = .2972). There was a main effect of drinking session (**Figure 1J**; *F*_drinking session_(3.461, 41.53) = 2.738; *p* = .0483) but no main effect of sex (*F*_sex_(1, 12) = 0.08197; *p* = .7795) nor an interaction (*F*_sex x drinking session_(11, 132) = 0.8800; *p* = .5616) on preference for 20% EtOH during adolescent IA2BC. There was not a sex difference in total EtOH consumption summed across all IA2BC sessions during adolescence (**Figure 1K**; *t*(12) = 0.4677, *p* = .6484).

In experiment 5, male and female Fisher 344 rats underwent a CIE paradigm starting at PND 31 (**Figure 1L-N**). A mixed-effects 2-way ANOVA (factors: drinking session, sex) indicated a main effect of drinking session on EtOH consumption (**Figure 1M**; *F*_drinking session_(23, 176) = 7.573; *p* < .0001) but not sex (*F*_sex_(1, 10) = 1.749; *p* = .2155) nor an interaction between drinking session and sex (*F*_sex x session_(23, 176) = 0.8760; *p* = .6303). Consistent with the adolescent IA2BC rat paradigm in Experiment 4, an unpaired t-test showed no sex difference in total EtOH consumption during adolescent CIE (**Figure 1N**; *t*(8) = 0.9178, *p* = .3855).

**Figure 1.**
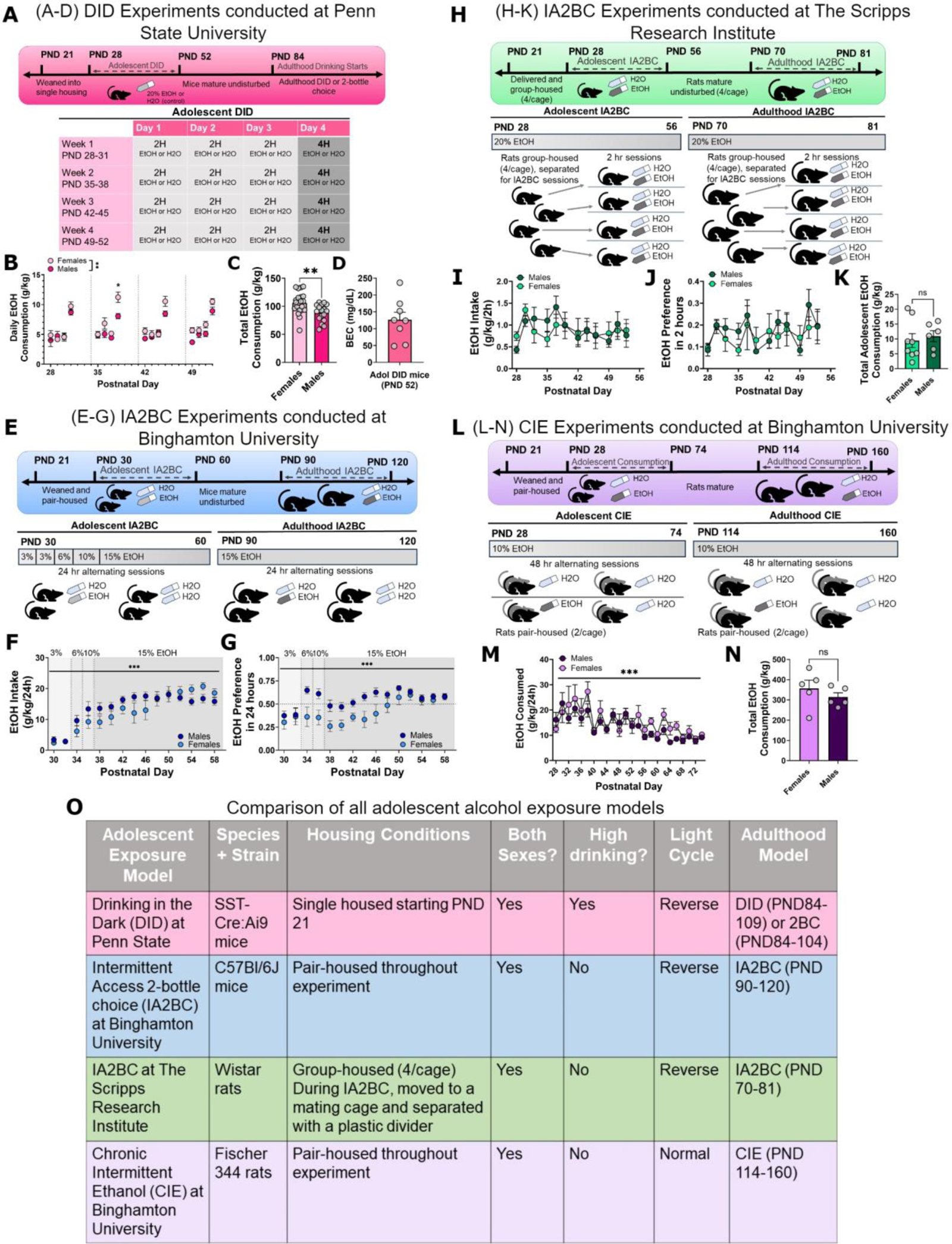
Overall methods and models. Experiments were conducted across three sites and four adolescent drinking paradigms. **A-D**, drinking in the dark model in mice, conducted at Penn State University; **E-G**, adolescent intermittent access model in mice, conducted at Binghamton University; **H-K** adolescent intermittent access model in rats, conducted at The Scripps Research Institute; and **L-N** adolescent chronic intermittent ethanol model in rats, conducted at Binghamton University. **O**, comparison of overall models. These four models cover a range of consumption paradigms and stress confounds such as paired or isolated housing. * = *p* < .05; ** = *p* < .01, *** = *p* < .001. PND: postnatal day; DID: drinking in the dark; IA2BC: intermittent access 2-bottle choice; CIE: chronic intermittent ethanol.

Overall, we were able to model adolescent alcohol exposure with several voluntary consumption paradigms across multiple housing conditions and in different model species (**Figure 1O**).

### 3.2 Sex and adolescent DID only have modest effects on adulthood binge drinking

To test changes in adulthood binge drinking due to adolescent binge alcohol exposure, mice were exposed to a “double DID” paradigm (timeline in **Figure 2A**). “Adol DID” mice underwent 4 weeks of DID during adolescence, while control mice had access to only water in adolescence. Beginning at PND 84, all mice underwent 4 weeks of adulthood DID. Alcohol consumption during the first 2-hour exposure, during the first 4-hour binge day, and across all 16 days of adulthood DID were analyzed using ANOVA and GLME (data from GLME are presented as Coefficient Estimate ± SEM).

A 2-way ANOVA revealed no effect of sex or adolescent DID on EtOH consumption during first adulthood 2 hr exposure (PND 84; **Figure 2B**; *F*_sex_(1, 38) = 0.1574, *p* = .6938, ω^2^ = 0.4077%; *F*_DID_(1, 38) = 0.08468, *p* = .7726, ω^2^ = 0.2194%; *F*_sex x DID_(1, 38) = 0.4489, *p* = .5069, ω^2^ = 1.163%). We found a significant main effect of sex on EtOH consumption at first adulthood binge day (**Figure 2C**; *F*_sex_(1, 42) = 6.098, *p* = .0177, ω^2^ = 12.17%), but no effect of adolescent DID nor a sex x DID interaction (*F*_DID_(1, 42) = 1.305, *p* = .2598, ω^2^ = 2.604%; *F*_sex x DID_(1, 42) = 0.9011, *p* = .3479, ω^2^ = 1.798%). During this first binge day, female mice consumed more EtOH than male mice. Although there were no significant effects of adolescent DID experience on daily adulthood EtOH drinking, there was a modest but significant main effect of adolescent DID on total EtOH consumption during adulthood DID (**Figure 2D**; *F*_DID_(1, 43) = 5.571, *p* = .0229, ω^2^ = 6.500%). There was also a main effect of sex on total EtOH consumption (*F*_sex_(1, 43) = 36.29, *p <* .0001, ω^2^ = 42.34%) but no interaction between sex and DID (*F*_sex x DID_(1, 43) = 0.2673, *p* = .6078, ω^2^ = 0.3119%). To confirm the mice drank to high BECs during adulthood DID, we collected blood samples via cardiac puncture within 30 minutes of the final adulthood binge session (PND 108; **Figure 2E**). Together, these results indicate that adolescent binge drinking only modestly increases total binge alcohol consumption in adulthood compared to mice without adolescent drinking experience.

Within the double DID mice, we assessed if changes in adulthood drinking are predicted by the amount of alcohol consumed during adolescence. We used a GLME with sex and total adolescent EtOH as predictor variables for each of the outcome variables of adulthood DID (consumption at first 2 hr exposure, first binge, and across all 16 DID sessions). The model accounted for litter as a random effect variable (**Table 1**; **Figure 2F-H**). Neither sex (**Figure 2F**; sex = male: 1.8215 ± 8.2076; *p =* .8274) nor total adolescent EtOH consumption (0.01726 ± 0.03444; *p* = 0.6236) were significant predictors of alcohol consumption during the first adulthood 2-hour exposure, nor was there an interaction between sex and total adolescent EtOH consumption (-0.02627 ± 0.0876; *p =* .7690). For EtOH consumption during first adulthood binge session, neither sex (**Figure 2G**; sex = male: -5.0827 ± 4.5009; *p =* .2736) nor total adolescent EtOH consumption (-0.01685 ± 0.02474; *p* = 0.5045) were significant predictors, nor was there an interaction between sex and total adolescent EtOH consumption (0.03506 ± 0.04712; *p =* .4664). Total adolescent EtOH consumption was a significant predictor for total EtOH consumption during adulthood DID (**Figure 2H**; 0.4907 ± 0.1687; *p* = .0090). Sex was not a significant predictor for total adulthood EtOH consumption (sex = male: -15.787 ± 30.926; *p* = .6156), nor was there an interaction (-0.03798 ± 0.3232, *p* = .9077).

**Table 1.**
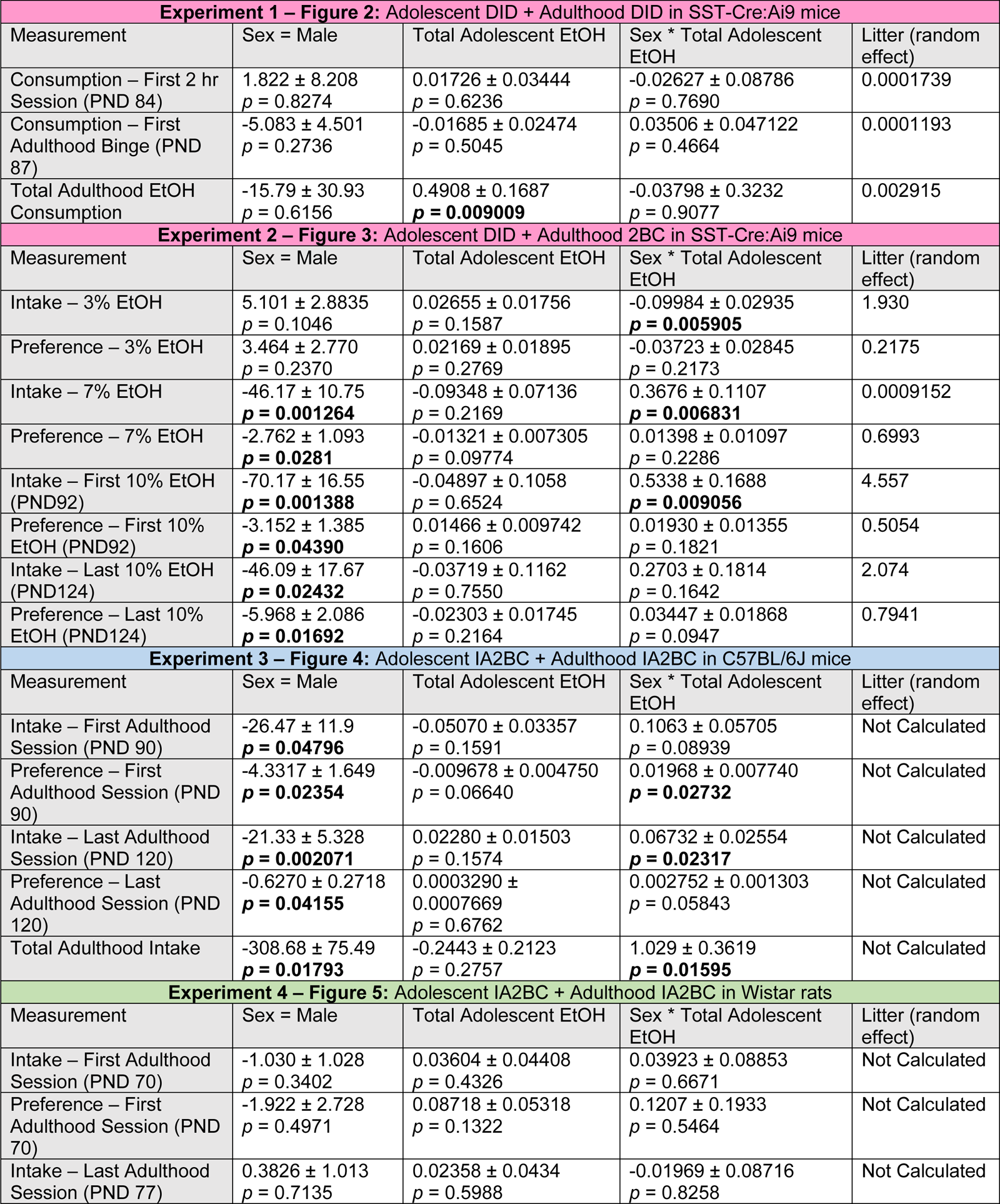
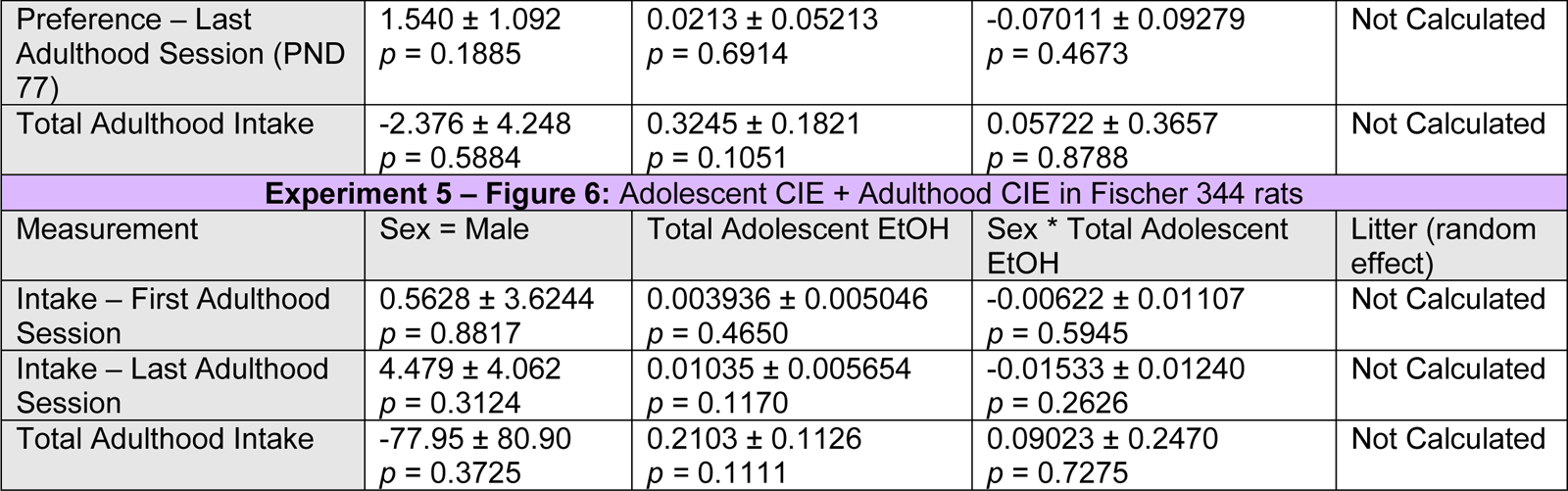
Results of GLME for all 5 adulthood drinking paradigms. Data are presented as the estimate of each fixed effect ± standard error. Significant predictors (*p* < .05) indicated in bold. DID: Drinking in the Dark; IA2BC: Intermittent Access 2-Bottle Choice; CIE: Chronic Intermittent Ethanol. Models for Experiments 1 and 2: [outcome variable] ∼ 1 + Sex*Total Adolescent EtOH + (1|litter) Models for Experiments 3, 4, 5: [outcome variable] ∼ 1 + Sex*Total Adolescent EtOH

**Experiment 1 – Figure 2: Adolescent DID + Adulthood DID in SST-Cre: Ai9 mice**
| Measurement | Sex = Male | Total Adolescent EtOH | Sex * Total Adolescent EtOH | Litter (random effect) |
| --- | --- | --- | --- | --- |
| Consumption – First 2 hr Session (PND 84) | 1.822 ± 8.208<br>*p* = 0.8274 | 0.01726 ± 0.03444<br>*p* = 0.6236 | -0.02627 ± 0.08786<br>*p* = 0.7690 | 0.0001739 |
| Consumption – First Adulthood Binge (PND 87) | -5.083 ± 4.501<br>*p* = 0.2736 | -0.01685 ± 0.02474<br>*p* = 0.5045 | 0.03506 ± 0.047122<br>*p* = 0.4664 | 0.0001193 |
| Total Adulthood EtOH Consumption | -15.79 ± 30.93<br>*p* = 0.6156 | 0.4908 ± 0.1687<br>***p* = 0.009009** | -0.03798 ± 0.3232<br>*p* = 0.9077 | 0.002915 |

**Experiment 2 – Figure 3: Adolescent DID + Adulthood 2BC in SST-Cre: Ai9 mice**
| Measurement | Sex = Male | Total Adolescent EtOH | Sex * Total Adolescent EtOH | Litter (random effect) |
| --- | --- | --- | --- | --- |
| Intake – 3% EtOH | 5.101 ± 2.8835<br>*p* = 0.1046 | 0.02655 ± 0.01756<br>*p* = 0.1587 | -0.09984 ± 0.02935<br>***p* = 0.005905** | 1.930 |
| Preference – 3% EtOH | 3.464 ± 2.770<br>*p* = 0.2370 | 0.02169 ± 0.01895<br>*p* = 0.2769 | -0.03723 ± 0.02845<br>*p* = 0.2173 | 0.2175 |
| Intake – 7% EtOH | -46.17 ± 10.75<br>***p* = 0.001264** | -0.09348 ± 0.07136<br>*p* = 0.2169 | 0.3676 ± 0.1107<br>***p* = 0.006831** | 0.0009152 |
| Preference – 7% EtOH | -2.762 ± 1.093<br>***p* = 0.0281** | -0.01321 ± 0.007305<br>*p* = 0.09774 | 0.01398 ± 0.01097<br>*p* = 0.2286 | 0.6993 |
| Intake – First 10% EtOH (PND92) | -70.17 ± 16.55<br>***p* = 0.001388** | -0.04897 ± 0.1058<br>*p* = 0.6524 | 0.5338 ± 0.1688<br>***p* = 0.009056** | 4.557 |
| Preference – First 10% EtOH (PND92) | -3.152 ± 1.385<br>***p* = 0.04390** | 0.01466 ± 0.009742<br>*p* = 0.1606 | 0.01930 ± 0.01355<br>*p* = 0.1821 | 0.5054 |
| Intake – Last 10% EtOH (PND124) | -46.09 ± 17.67<br>***p* = 0.02432** | -0.03719 ± 0.1162<br>*p* = 0.7550 | 0.2703 ± 0.1814<br>*p* = 0.1642 | 2.074 |
| Preference – Last 10% EtOH (PND124) | -5.968 ± 2.086<br>***p* = 0.01692** | -0.02303 ± 0.01745<br>*p* = 0.2164 | 0.03447 ± 0.01868<br>*p* = 0.0947 | 0.7941 |

**Experiment 3 – Figure 4: Adolescent IA2BC + Adulthood IA2BC in C57BL/6J mice**
| Measurement | Sex = Male | Total Adolescent EtOH | Sex * Total Adolescent EtOH | Litter (random effect) |
| --- | --- | --- | --- | --- |
| Intake – First Adulthood Session (PND 90) | -26.47 ± 11.9<br>***p* = 0.04796** | -0.05070 ± 0.03357<br>*p* = 0.1591 | 0.1063 ± 0.05705<br>*p* = 0.08939 | Not Calculated |
| Preference – First Adulthood Session (PND 90) | -4.3317 ± 1.649<br>***p* = 0.02354** | -0.009678 ± 0.004750<br>*p* = 0.06640 | 0.01968 ± 0.007740<br>***p* = 0.02732** | Not Calculated |
| Intake – Last Adulthood Session (PND 120) | -21.33 ± 5.328<br>***p* = 0.002071** | 0.02280 ± 0.01503<br>*p* = 0.1574 | 0.06732 ± 0.02554<br>***p* = 0.02317** | Not Calculated |
| Preference – Last Adulthood Session (PND 120) | -0.6270 ± 0.2718<br>***p* = 0.04155** | 0.0003290 ± 0.0007669<br>*p* = 0.6762 | 0.002752 ± 0.001303<br>*p* = 0.05843 | Not Calculated |
| Total Adulthood Intake | -308.68 ± 75.49<br>***p* = 0.01793** | -0.2443 ± 0.2123<br>*p* = 0.2757 | 1.029 ± 0.3619<br>***p* = 0.01595** | Not Calculated |

**Table 1.** Results of GLME for all 5 adulthood drinking paradigms.
| Measurement | Sex = Male | Total Adolescent EtOH | Sex * Total Adolescent EtOH | Litter (random effect) |
| --- | --- | --- | --- | --- |
| Intake – First Adulthood Session (PND 70) | -1.030 ± 1.028<br>*p* = 0.3402 | 0.03604 ± 0.04408<br>*p* = 0.4326 | 0.03923 ± 0.08853<br>*p* = 0.6671 | Not Calculated |
| Preference – First Adulthood Session (PND 70) | -1.922 ± 2.728<br>*p* = 0.4971 | 0.08718 ± 0.05318<br>*p* = 0.1322 | 0.1207 ± 0.1933<br>*p* = 0.5464 | Not Calculated |
| Intake – Last Adulthood Session (PND 77) | 0.3826 ± 1.013<br>*p* = 0.7135 | 0.02358 ± 0.0434<br>*p* = 0.5988 | -0.01969 ± 0.08716<br>*p* = 0.8258 | Not Calculated |
| Preference – Last Adulthood Session (PND 77) | 1.540 ± 1.092<br>*p* = 0.1885 | 0.0213 ± 0.05213<br>*p* = 0.6914 | -0.07011 ± 0.09279<br>*p* = 0.4673 | Not Calculated |
| Total Adulthood Intake | -2.376 ± 4.248<br>*p* = 0.5884 | 0.3245 ± 0.1821<br>*p* = 0.1051 | 0.05722 ± 0.3657<br>*p* = 0.8788 | Not Calculated |
| **Experiment 5 – Figure 6: Adolescent CIE + Adulthood CIE in Fischer 344 rats** |  |  |  |  |
| Measurement | Sex = Male | Total Adolescent EtOH | Sex * Total Adolescent EtOH | Litter (random effect) |
| Intake – First Adulthood Session | 0.5628 ± 3.6244<br>*p* = 0.8817 | 0.003936 ± 0.005046<br>*p* = 0.4650 | -0.00622 ± 0.01107<br>*p* = 0.5945 | Not Calculated |
| Intake – Last Adulthood Session | 4.479 ± 4.062<br>*p* = 0.3124 | 0.01035 ± 0.005654<br>*p* = 0.1170 | -0.01533 ± 0.01240<br>*p* = 0.2626 | Not Calculated |
| Total Adulthood Intake | -77.95 ± 80.90<br>*p* = 0.3725 | 0.2103 ± 0.1126<br>*p* = 0.1111 | 0.09023 ± 0.2470<br>*p* = 0.7275 | Not Calculated |

We used a mixed-effects 3-way ANOVA (factors: binge day, adolescent DID, sex) to analyze changes in binge drinking throughout 4 cycles of adulthood DID (daily EtOH consumption for all 16 days of adulthood DID shown in **Figure 2I**). We found significant main effects of sex (*F*_sex_(1, 43) = 26.69, *p* < .0001), binge day (*F*_binge_(3, 126) = 2.798, *p* = .0429), and adolescent DID history (*F*_DID_(1, 43) = 4.198, *p* = .0466). All interactions were not significant (*p* > .05). Mice were weighed weekly to assess whether adolescent binge drinking affected physical maturation (**Figure 2J**). A mixed-effects 3-way ANOVA (factors: age, adolescent DID, sex) revealed an expected interaction between age and sex (*F*_age x DID_(11, 445) = 9.655, *p* < .0001), but no main effect of adolescent DID (*F*_DID_(1, 43) = 0.2329, *p* = .6318; all other interactions not significant).

**Figure 2.**
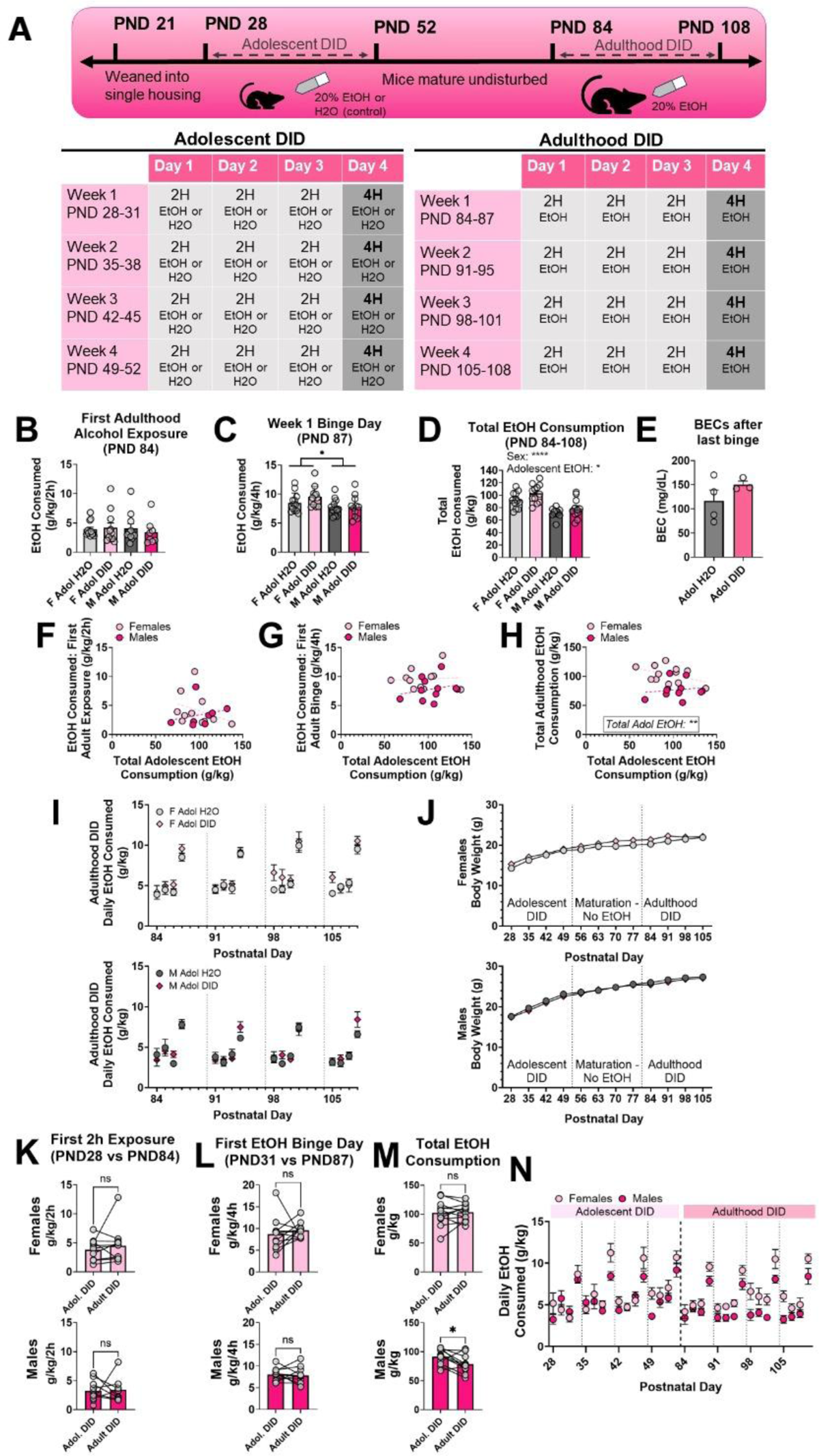
Sex and total adolescent alcohol consumption only modestly increase total alcohol consumption during adulthood DID in mice. **(A)** Experimental timeline. **(B)** Neither Sex nor adolescent DID exposure affect EtOH consumption on first adulthood 2-hour exposure at PND 84. **(C)** Female mice drank more than male mice during the first 4-hour binge session at PND 87, but this effect was not modulated by adolescent DID. **(D)** There were main effects of sex and adolescent DID on total EtOH consumption across all 16 days of adulthood DID. **(E)** Adolescent DID experience did not affect BECs achieved during the final adulthood binge session (*n* = 4 Adol H2O mice, *n =* 3 Adol DID mice, sexes combined). **(F-H)** Relationships between total adolescent EtOH consumption and each of the 3 adult binge drinking parameters – EtOH consumed in first 2-hour exposure, first 4-hour binge, and across all 16 sessions of DID. Significant predictors identified by GLME are indicated where appropriate. **(I)** Daily EtOH consumption during adulthood DID in female (top) and male (bottom) mice. **(J)** Adolescent DID did not affect body weight in female (top) or male (bottom) mice. **(K)** Female and male Adolescent DID mice do not change their drinking during the first 2-hour exposure in adolescence (PND 28) compared to in adulthood (PND 84). **(L)** There was no difference in EtOH consumption during first binge day in adolescence (PND 31) compared to adulthood (PND 87). **(M)** While female mice do not show changes in total EtOH consumption in adolescence and adulthood DID, male mice show reduced total EtOH intake during adulthood DID. **(N)** Daily EtOH consumption in adolescent and adulthood DID in double drinking mice. Trendlines in F-H represent the line of best fit within each sex. Significant main effects are indicated as * = *p* < .05, **** = *p* < .0001

We also assessed whether mice exposed to adolescent DID showed any changes in their drinking patterns from adolescence to adulthood. We used paired t-tests to compare EtOH consumption at first 2-hour exposure, first 4-hour binge day, and across all 16 days of DID in adolescence and adulthood. There was no change in EtOH consumption during first 2-hour exposures (PND 28 vs PND 84) in female (**Figure 2K**; *t*(10) = 0.8384, *p* = .4214) or male (*t*(7) = 0.1440, *p* = .8896) double drinking mice. Similarly, neither female (**Figure 2L**; *t*(10) = 0.6363, *p* = .5389) nor male mice (*t*(10) = 0.2803, *p* = .7850) showed differences in drinking at first binge day (PND 31 vs PND 87). Although there was no difference in total EtOH consumption during adolescence and adult DID in female mice (**Figure 2M-N**; *t*(11) = 0.2006, *p* = .8447), male mice drank less total EtOH, normalized to body weight, during adulthood DID than during adolescent DID (*t*(10) = 2.607, *p* = .0262). These within-subject results indicate that mice exposed to adolescent DID did not increase their ethanol consumption during adulthood binge drinking. Taken together, these findings indicate that mice exposed to adolescent DID show only a modest increase in total EtOH binge consumption compared to mice not exposed to EtOH until adulthood.

### 3.3 Sex and Adolescent DID Effects on Adulthood Ethanol Preference

To determine whether adolescent DID alters preference for alcohol in adulthood, a separate cohort of SST-Cre:Ai9 mice underwent DID or water control during adolescence and then 6 weeks of 2-bottle choice during adulthood beginning PND 84 (**Figure 3A**).

We analyzed the effects of adolescent drinking and sex on EtOH intake and preference on 4 sessions of the 2BC procedure: the 3% EtOH session (PND 84), the final 7% EtOH session (PND 90), the first 10% EtOH session (PND 92) and the final 10% EtOH session (PND 124). We used 2-way ANOVA and GLMEs to assess the effects of sex and total adolescent alcohol consumption on each of the outcome variables.

**Figure 3.**
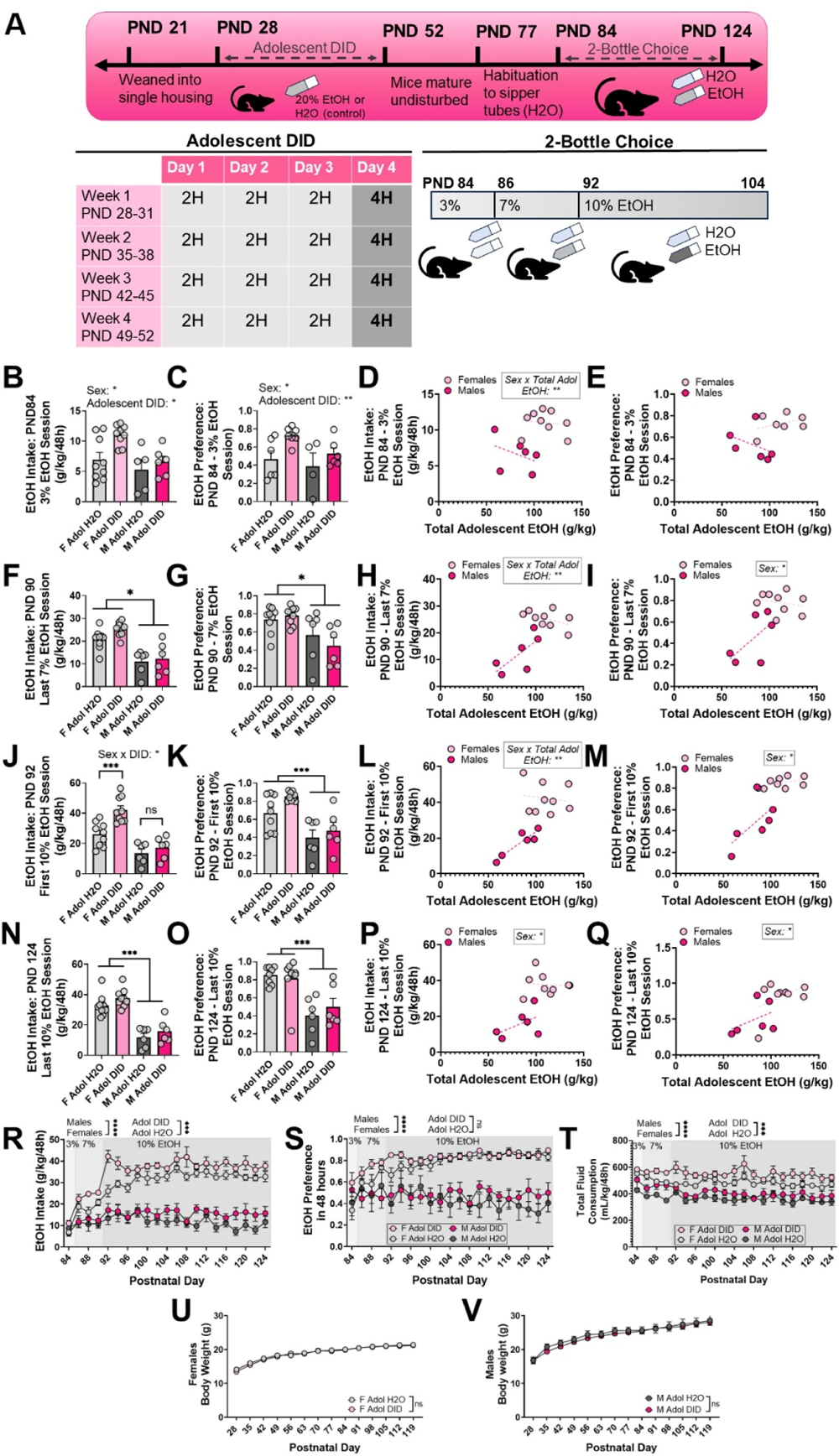
Intake and preference for EtOH during a 2-bottle choice model after adolescent binge drinking in SST-Cre:Ai9 mice. (A) Experimental timeline. (B-E) Mice which went through adolescent DID show increased preference and intake for 3% EtOH on PND 84. (F-I) Female mice show a higher preference and intake of 7% EtOH on PND 90. (J-M) Preference and intake of 10% EtOH are highest in female mice which underwent adolescent DID. (N-Q) Sex influences both preference and intake of 10% EtOH at PND 104. (R-T) Daily EtOH intake, preference, and total fluid consumption during adulthood 2-bottle choice. (U-V) Body weight is not affected by adolescent DID. Trendlines in D-E, H-I, L-M, P-Q indicate the lines of best fit for each sex. * = *p* < .05, ** = *p* < .01, *** = *p* < .001, **** = *p* < .0001.

During the first adulthood 2BC session, a 2-way ANOVA (factors: sex, adolescent DID) found main effects of both sex (*F*_sex_(1, 25) = 7.755, *p* = .0101, ω^2^ = 18.29%) and adolescent DID (**Figure 3B**; *F*_DID_(1, 25) = 6.001, *p* = .0216, ω^2^ = 14.15%) on 3% EtOH intake, but no interaction (*F*_sex x DID_(1, 25) = 1.715, *p* = .2022, ω^2^ = 4.046%). There was a significant main effect of DID on preference for 3% EtOH (**Figure 3C**; *F*_DID_(1, 20) = 8.393, *p* = .0089, ω^2^ = 25.04%) and an effect of sex (*F*_sex_(1, 20) = 4.533, *p* = .7339, ω^2^ = 13.52%) with no interaction (*F*_sex x DID_(1, 20) = 0.2046, *p* = .6559, ω^2^ = 0.6104%). A GLME accounting for sex and total adolescent EtOH intake within the adolescent DID mice indicated that the interaction between sex and total adolescent EtOH consumption was a significant predictor for intake during the 3% EtOH session (GLM results in **Table 1**; **Figure 3D**; -0.09984 ± 0.02935, *p =* .0059) but not sex (5.1013 ± 2.8835, *p =* .1046) or total adolescent EtOH separately (0.02655 ± 0.01756, *p* = .1587). Similarly, for preference during the 3% EtOH session, the interaction between sex and total adolescent EtOH was a significant predictor (**Figure 3E**; -0.03719 ± 0.01011, *p* = .2173) but not sex (2.209 ± 1.0267, *p* = .0600) nor total adolescent EtOH (0.01288 ± 0.006588, *p* = .0823).

During the last 7% EtOH session on PND 90, there was a significant effect of sex on EtOH intake during the last 7% session (**Figure 3F**; *F*_sex_(1, 26) = 40.31, *p* < .0001, ω^2^ = 57.20%) but not adolescent DID (*F*_DID_(1, 26) = 2.647, *p* = .1158, ω^2^ = 3.757%) nor an interaction (*F*_sex x DID_(1, 26) = 0.7730, *p* = .3874, ω^2^ = 1.097%). There was a main effect of sex (**Figure 3G**; *F*_sex_(1, 26) = 12.90, *p* = .0013, ω^2^ = 31.97%) but not adolescent DID (*F*_DID_(1, 26) = 0.2834, *p* = .5990, ω^2^ = 0.7026%) nor an interaction (*F*_sex x DID_(1, 26) = 1.344, *p* = .2569, ω^2^ = 3.332%) on preference for 7% EtOH. Within the mice which drank EtOH during adolescence, the interaction between sex and total adolescent EtOH was a significant predictor of EtOH intake during the PND 90 session (**Figure 3H**; 0.3676 ± .1107, *p* = .006831). Sex (**Figure 3I**; sex = male: -2.762 ± 1.0928, *p* = .02811) but not total adolescent EtOH consumption (-0.01322 ± 0.007305, *p* = .09774) nor an interaction (0.01398 ± 0.01097, *p* = .2286) was a significant predictor of preference for EtOH during the final 7% EtOH session.

For intake at the first 10% drinking session, there was a significant interaction between sex and adolescent DID (**Figure 3J**; *F*_sex x DID_(1, 26) = 4.390, *p* = .0460, ω^2^ = 5.097%). Fisher’s post-hoc comparisons revealed that within the female mice, the mice which underwent adolescent DID consumed significantly more EtOH than water-exposed female mice (*p* = .0002), while there was no effect of adolescent DID in the male mice (*p* = .4160). At the first 10% drinking session on PND 92, there was a main effect of sex (**Figure 3K**; *F*_sex_(1, 26) = 24.27, *p* < .0001, ω^2^ = 43.33%) but not adolescent DID (*F*_DID_(1, 26) = 4.159, *p* = .0517, ω^2^ = 7.426%) nor an interaction (*F*_sex x DID_(1, 26) = 0.6797, *p* = .4172, ω^2^ = 1.214%). Within the mice that were exposed to adolescent DID, the interaction between sex and total adolescent EtOH intake was a significant predictor of EtOH intake during the first 10% drinking session (**Figure 3L**; 0.5338 ± 0.16884, *p* = .009056). Sex was a significant predictor of preference during the first 10% drinking session (**Figure 3M**; sex = male: -3.1521 ± 1.3854, *p* = .04390) but not total adolescent EtOH (0.01466 ± 0.009742, *p* = .1606) or an interaction (0.1930 ± 0.01355, *p* = .1821).

At the final 10% drinking session on PND 124, there was a main effect of sex on EtOH intake (**Figure 3N**; *F*_sex_(1, 26) = 71.15, *p* < .0001, ω^2^ = 70.76%) but no effect of adolescent DID (*F*_DID_(1, 26) = 3.109, *p* = .0896, ω^2^ = 3.092%) nor an interaction (*F*_sex x DID_(1, 26) = 0.03309, *p =* .8571, ω^2^ = 0.03290%). There was a main effect of sex on preference for EtOH during this final session (**Figure 3O**; *F*_sex_(1, 25) = 26.73, *p* < .0001, ω^2^ = 50.97%) but not adolescent DID (*F*_DID_(1, 25) = 0.1854, *p =* .6705, ω^2^ = 0.3535%) or an interaction (*F*_sex x DID_(1, 25) = 0.7128, *p* = .4065, ω^2^ = 1.359%). Within the mice exposed to adolescent DID, sex was a significant predictor of intake during the final 10% session (**Figure 3P**; sex = male: -46.09 ± 17.67, *p* = .02432), but not total adolescent EtOH (-0.03719 ± 0.1162, *p* = .7550) nor the interaction (0.2703 ± 0.1814, *p* = .1642). Sex was a significant predictor of preference during the final 10% EtOH session (**Figure 3Q**; sex = male: -5.978 ± 2.086, *p* = .01692) but not total adolescent EtOH (-.02303 ± 0.01745, *p* = .2164) nor an interaction (0.03447 ± 0.01868, *p* = .09473).

To get a broader understanding of changes in EtOH intake and preference throughout 6 weeks of drinking, we analyzed the area under the curve (AUC) of the daily drinking trends (**Figure 3R-T**). A 2-way ANOVA (factors: sex, adolescent DID) revealed significant main effects of sex (**Figure 3R**; *F*_sex_(1, 26) = 304.0, *p* < .0001) and adolescent DID (*F*_DID_(1, 26) = 17.21, *p* = .0003) on EtOH intake AUC but not an interaction between sex and adolescent DID (*F*_sex x DID_(1, 26) = 1.263, *p* = .2713). Šídák’s multiple comparisons test indicated that adolescent DID significantly increased EtOH intake in female (adjusted *p* = .0006) but not male mice (adjusted *p* = .1197). For EtOH preference AUC, there was a main effect of sex (**Figure 3S**; *F*_sex_(1, 26) = 162.3, *p* < .0001) but not adolescent DID (*F*_DID_(1, 26) = 2.974, *p* = .0965) nor an interaction (*F*_sex x DID_(1, 26) = 0.005052, *p* = .9439). Because adolescent DID experience had a significant effect on EtOH intake but not preference during adulthood 2BC, we also looked at changes in total fluid consumption (H2O + EtOH; mL/kg) during adulthood 2BC (**Figure 3T**). There were significant effects of sex (*F*_sex_(1, 26) = 111.2, *p* < .0001) and adolescent DID (*F*_DID_(1, 26) = 19.88, *p* = .0001) on AUC for total fluid consumption, but no interaction (*F*_sex x DID_(1, 26) = 0.8252, *p* = .3720). Adolescent DID increased total fluid consumption during adulthood 2BC without altering preference for EtOH, indicating that our observed increase in EtOH intake is not specific to EtOH.

A 3-way mixed-effects ANOVA (factors: sex, adolescent DID, age) revealed a significant interaction between sex and age on body weight (**Figure 3U-V**; *F*_sex x age_(13, 283) = 17.33, *p* < .0001) but no effect of adolescent DID (*F*_DID_(1, 26) = 0.4205, *p* = .5224; all other interactions *p* > .05). Neither binge drinking during adolescence nor 2-bottle choice in adulthood affected the physical growth of SST-Cre:Ai9 mice. Together, this experiment identified modest changes in EtOH preference and intake in mice exposed to adolescent DID, but these changes did not persist through the end of the 2BC protocol.

### 3.4 Sex and Adolescent Alcohol Effects on Drinking in a Mouse Intermittent Access 2-bottle choice Procedure Conducted at Binghamton University

In Experiment 3, changes in adulthood EtOH consumption were investigated in an Intermittent Access 2-bottle choice (IA2BC) protocol in pair-housed adolescent and adult C57BL/6J mice (**Figure 4A**).

A 2-way ANOVA revealed that intake of 15% EtOH during the first adulthood 2BC session on PND 90 was not significantly changed as a function of sex (**Figure 4B**; *F*_sex_(1, 27) = 3.707, *p* = .0648, ω^2^ = 11.87%), adolescent EtOH exposure (*F*_Adol EtOH_(1, 27) = 0.5876, *p* = .4500, ω^2^ = 1.882%), or an interaction (*F*_sex x Adol EtOH_(1, 27) = 0.2024, *p* = .6564, ω^2^ = 0.6483%). There was no main effect of sex (**Figure 4C**; *F*_sex_(1, 27) = 0.7574, *p* = .3918, ω^2^ = 2.629%), adolescent EtOH exposure (*F*_Adol EtOH_(1, 27) = 1.116, *p* = .3001, ω^2^ = 3.874%), or an interaction (*F*_sex x Adol EtOH_(1, 27) = 0.09315, *p* = .7626, ω^2^ = 0.3233%) on preference for 15% EtOH at the first adulthood drinking session. Within the mice which underwent adolescent IA2BC, sex was a significant predictor of EtOH intake during the first adulthood drinking session (GLME results in **Table 1**; **Figure 4D**; sex = male: - 26.474 ± 11.9, *p* = .04796) but not total adolescent EtOH intake (-0.05070 ± 0.03357, *p* = .1591) nor the interaction between sex and total adolescent EtOH intake (0.1063 ± 0.05705, *p* = .0894). The interaction between sex and total adolescent EtOH intake was a significant predictor of EtOH preference during the first adulthood IA2BC session (**Figure 4E**; 0.019683 ± 0.007740, *p* = .02732), but not total adolescent EtOH intake (-0.009678 ± 0.004750, *p* = .06640).

There was a significant main effect of sex on intake during the last day of adulthood IA2BC on PND 120 (**Figure 4F**; *F*_sex_(1, 27) = 10.38, *p* = .0033, ω^2^ = 25.52%) but not adolescent EtOH exposure (*F*_Adol EtOH_(1, 27) = 0.5118, *p* = .4805, ω^2^ = 1.258%) nor an interaction (*F*_sex x Adol EtOH_(1, 27) = 2.603, *p* = .1183, ω^2^ = 6.398%). Preference for EtOH during the last IA2BC session was not significantly affected by sex (**Figure 4G**; *F*_sex_(1, 27) = 0.9153, *p* = .3472, ω^2^ = 2.810%), adolescent EtOH exposure (*F*_Adol EtOH_(1, 27) = 0.7364, *p* = .3984, ω^2^ = 2.261%), or an interaction (*F*_sex x Adol EtOH_(1, 27) = 4.046, *p* = .0544, ω^2^ = 12.42%). The interaction between sex and total adolescent EtOH intake was a significant predictor of EtOH intake during the last adulthood drinking session (**Figure 4H**; 0.067321 ± 0.02554, *p* = .02317). Sex was a significant predictor of EtOH preference during the last drinking session (**Figure 4I**; sex = male: -0.6270 ± 0.2718, *p =* .04155) but not total adolescent EtOH intake (0.0003290 ± 0.2718, *p* = .6762) or the interaction between sex and total adolescent EtOH intake (0.002752 ± 0.001303, *p* = .0584).

**Figure 4.**
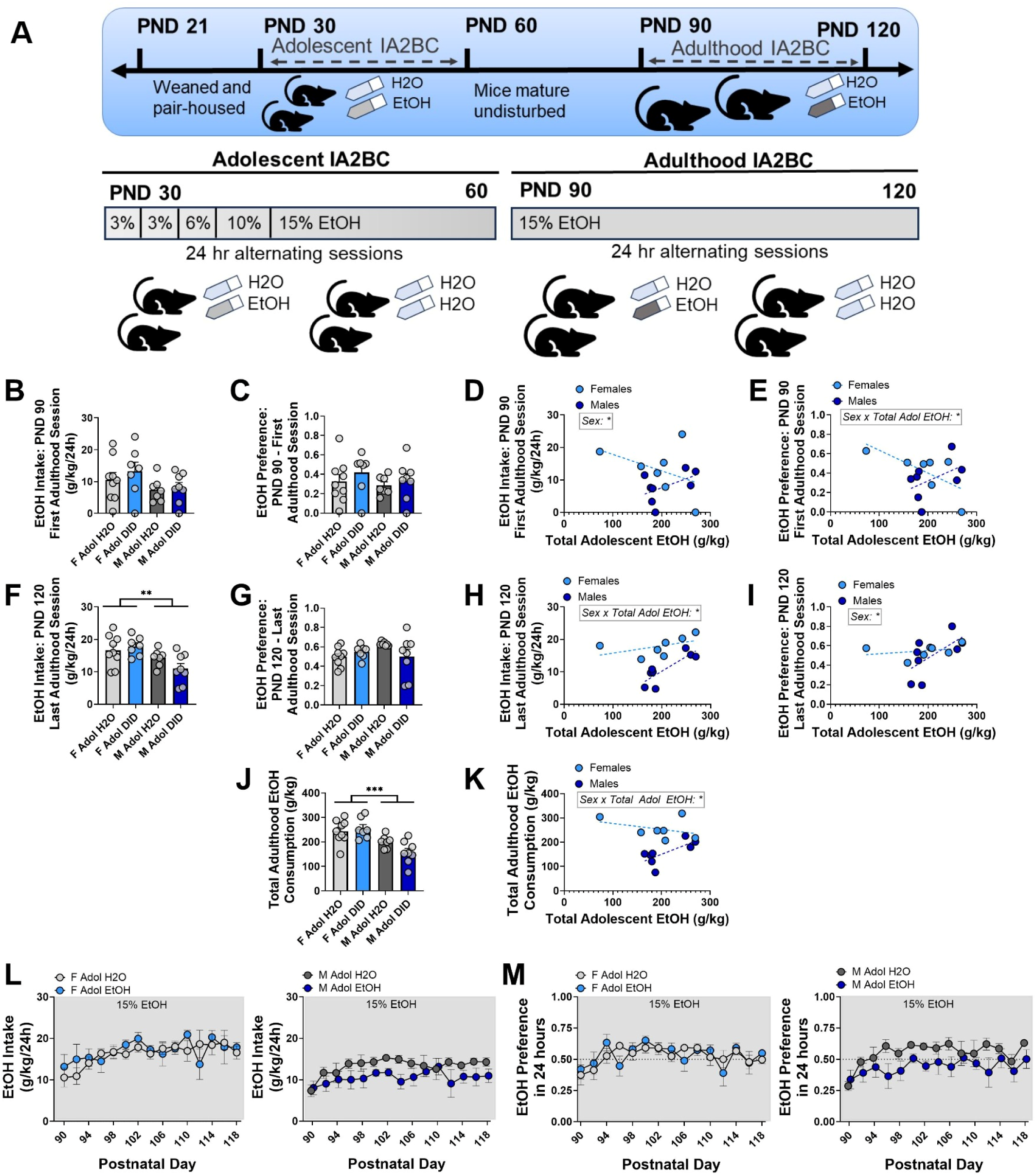
Pair-housed C57Bl/6J mice do not show changes in adulthood EtOH consumption in an IA2BC model. (A) Experimental schematic. (B-E) Neither EtOH preference nor intake was altered as a function of sex or adolescent EtOH exposure during the first adulthood IA2BC session on PND 90. (F-I) Female mice consumed significantly more EtOH during the final adulthood IA2BC session on PND 120, but this was not modulated by adolescent EtOH exposure. (J-K) Female mice consumed more EtOH across all 15 adulthood drinking sessions. (L-M) EtOH intake and preference across all 15 IA2BC sessions, split by sex. Significant predictors identified by GLME are indicated on each graph. Trendlines in D-E, H-I, K represent the line of best fit for each sex. B-K: Each value represents one cage with two mice. ** = *p* < .01, *** = *p* < .001.

Total adulthood EtOH intake across all 15 IA2BC sessions in adulthood was significantly affected by sex (**Figure 4J**; *F*_sex_(1, 27) = 21.81, *p* < .0001, ω^2^ = 40.58%), but not adolescent EtOH exposure (*F*_Adol EtOH_(1, 27) = 1.158, *p* = .2914, ω^2^ = 2.154%) or an interaction (*F*_sex x Adol EtOH_(1, 27) = 3.180, *p* = .0858, ω^2^ = 5.916%). Within the mice exposed to EtOH in adolescence, the interaction between sex and total adolescent EtOH intake was a significant predictor of total adulthood intake (**Figure 4K**; 1.0294 ± 0.3619, *p* = .01600). In adulthood, female pair-housed C57BL/6J mice consumed more EtOH across IA2BC sessions compared to male mice, but there was not a robust increase in EtOH intake in adult mice which were exposed to EtOH in adolescence compared to their adolescent water counterparts (daily intake and preference shown in **Figure 4L-M**). Together, this experiment did not show reliable escalations in EtOH intake or preference in mice exposed to adolescent EtOH.

### 3.5 Sex and Adolescent Alcohol Effects on Drinking in a Rat Intermittent Access 2-bottle Choice Procedure Conducted at Scripps

Adult (PND 70) Wistar rats underwent 6 sessions of IA2BC following 2 weeks of abstinence after the conclusion of adolescent IA2BC (**Figure 5**). As in adolescence, rats were group-housed and separated with a divider for 2-hour sessions of 2-bottle choice every other day (i.e. Tuesday, Thursday, Saturday; timeline in **Figure 5A**, daily EtOH intake and preference in **Figure 5B-C**). This experiment allowed us to assess changes in adulthood drinking within rats exposed to EtOH in adolescence. A 2-way repeated-measures ANOVA (factors: sex, drinking session – PND 56 and PND 70) indicated no effect of drinking session (**Figure 5D**; *F*_drinking session_(1, 12) = 0.7056, *p* = .4173, ω^2^ = 1.670%), sex (*F*_sex_(1, 12) = 0.8807, *p* = .3665, ω^2^ = 4.611%), nor any interaction (*F*_sex x drinking session_(1, 12) = 0.800, *p* = .3887, ω^2^ = 1.894%) on EtOH intake. Rats exposed to EtOH in adolescence did not increase their EtOH intake in adulthood.

**Figure 5.**
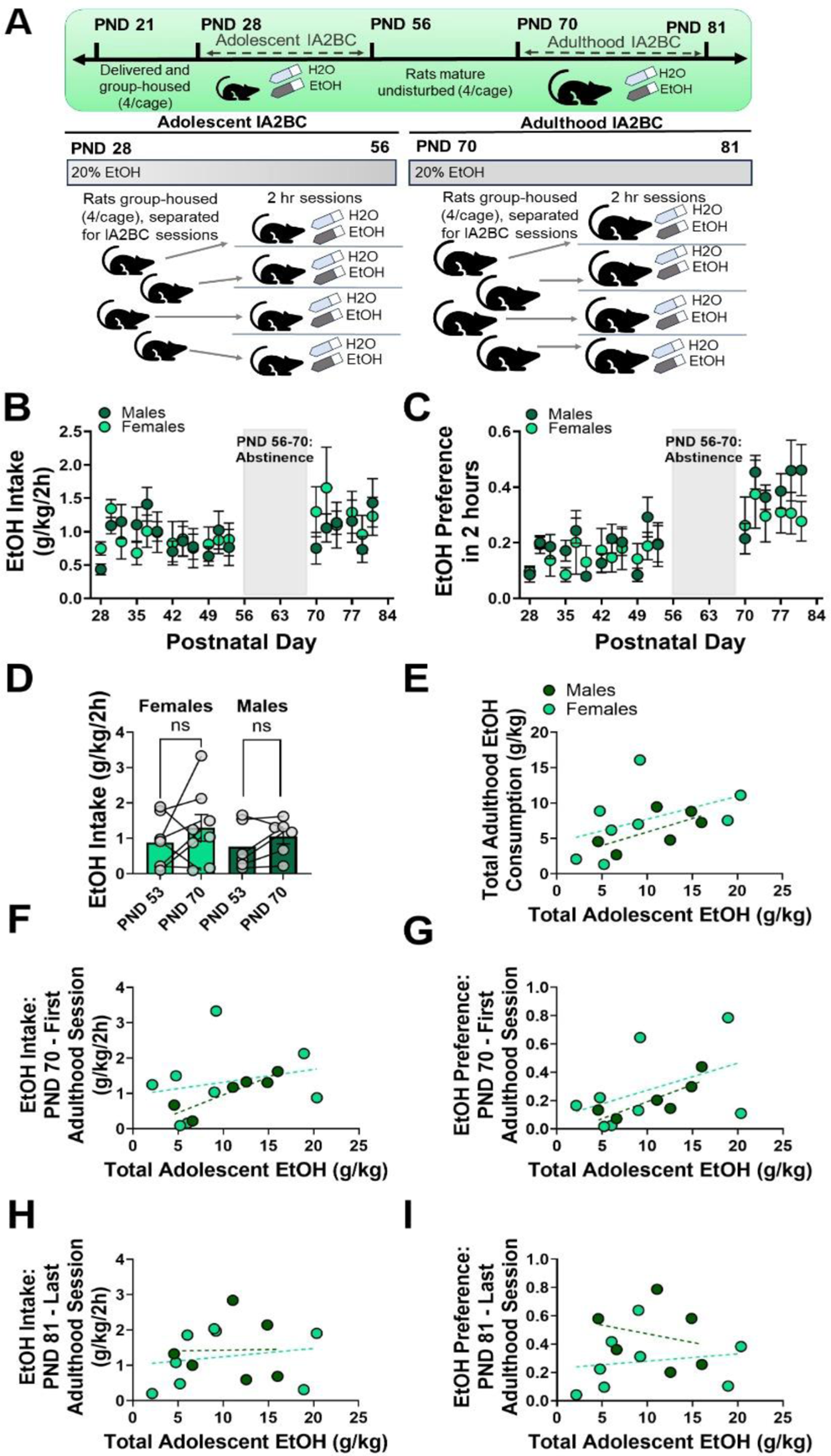
Group housed Wistar rats do not show increases in adulthood IA2BC following adolescent IA2BC. **(A)** Experimental timeline. **(B – C)** Rats that underwent adolescent alcohol exposure do not show increased EtOH intake or preference during adulthood. **(D)** There is no significant effect of drinking session on EtOH intake following adolescent alcohol exposure. **(E – I)** Total adolescent EtOH consumption is not a significant predictor of total adulthood EtOH consumption, first adulthood EtOH exposure intake or preference, or last adulthood EtOH exposure intake or preference. Trendlines in E-I represent the line of best fit for each sex.

As with our other models, we used a GLME to determine whether sex or total adolescent EtOH consumption are significant predictors of measurements of adulthood EtOH consumption. For this IA2BC experiment, the measurements of adulthood EtOH consumption were: total EtOH intake, EtOH preference during the first adulthood session (PND 70), intake during the first adulthood session (PND 70), preference during the final adulthood session (PND 81), and intake during the final adulthood session (PND 81) across the 6 sessions of IA2BC (GLME results in **Table 1**). Total EtOH intake across 6 sessions of adulthood IA2BC was not predicted by sex (**Figure 5E**; -2.3755 ± 4.2482, *p* = .5884) or total adolescent EtOH (0.32447 ± 0.18209, *p =* .1051), nor an interaction (0.057217 ± 0.3657, *p* = .8788). Neither sex (**Figure 5F**; -1.9221 ± 2.7278, *p* = .4971) nor total adolescent EtOH (0.08718 ± 0.05318, *p* = .1322) were significant predictors of EtOH preference during the first adulthood drinking session, nor was there an interaction (0.1207 ± 0.1933, *p* = .5464). There was no effect of sex (**Figure 5G**; -1.03 ± 1.0284, *p* = .3402) or total adolescent EtOH (0.03604 ± 0.04408, *p* = .4326), nor an interaction (0.03923 ± 0.08853, *p* = .6671) on EtOH intake during the first adulthood drinking session. For the final drinking session, preference was not significantly predicted by sex (**Figure 5H**; 1.5404 ± 1.0915, *p* = .1885), total adolescent EtOH (0.0213 ± 0.05213, *p =* .6914), or an interaction between sex and total adolescent EtOH (-0.07011 ± 0.09279, *p* = .4673). EtOH intake during this final session was not predicted by sex (**Figure 5I**; 0.38257 ± 1.0125, *p* = .7135), total adolescent EtOH (0.02358, *p* = .5988), or an interaction (-0.01969 ± 0.08716, *p* = .8258).

### 3.6 Adolescent Chronic Intermittent Ethanol Does Not Increase Adulthood Alcohol Consumption

Adult male and female Fischer 344 rats underwent a CIE paradigm of EtOH consumption beginning at PND 114 (**Figure 6**, experimental timeline in **Figure 6A**). This experiment allowed us to investigate any changes in adulthood drinking phenotypes within rats exposed to adolescent EtOH. A mixed-effects ANOVA indicated a main effect of time on EtOH intake, such that rats consumed less EtOH across time during adolescence and into adulthood (**Figure 6B**; *F_session_*(4.335, 33.85) = 14.07; *p* < .0001). There was also a main effect of sex on EtOH intake, where females consumed more EtOH than males (*F*_sex_(1, 10) = 6.296; *p* = .0310); however there was no significant interaction between sex and age of drinking session (*F*_sex x session_(47, 367) = 0.9696; *p* = .5333). Paired t-tests revealed that rats consumed less total EtOH across all twelve cycles of CIE in adulthood than in adolescence, in both females (**Figure 6C**; *t*(4) = 9.329, *p* = .0007) and males (*t*(4) = 3.884, *p* = .0178).

GLME was used to examine the interaction between total adolescent ethanol consumption and sex on measurements of adulthood drinking, including EtOH intake during the first adulthood CIE session, EtOH during the final adulthood CIE session, and total EtOH intake during adulthood CIE. Sex (**Figure 6E**; sex = male: 0.5628 ± 3.6244, *p =* .8817) nor total adolescent EtOH intake (0.003936 ± 0.005046, *p =* .4650), nor the interaction (-0.00622 ± 0.01107, *p =* .5945), were significant predictors of intake during the first adulthood CIE session. There were no significant predictors of EtOH intake during the last CIE session, including (**Figure 6F**; sex = male: 4.4785 ± 4.0616; *p* = .3124), total adolescent EtOH (0.01035 ± 0.005654, *p* = .1170), nor the interaction (-0.01533 ± 0.01240, *p* = .2626). Finally, sex (**Figure 6G**; sex = male: -77.953 ± 80.901, *p* = .3725), total adolescent EtOH (0.2103 ± 0.1126, *p* = .1111), nor the interaction (0.09023 ± 0.2470, *p* = 0.7275) were significant predictors of total EtOH intake during adulthood drinking sessions. Together, the results from both models in rats indicate that rats exposed to alcohol in adolescence do not increase their drinking in adulthood.

**Figure 6.**
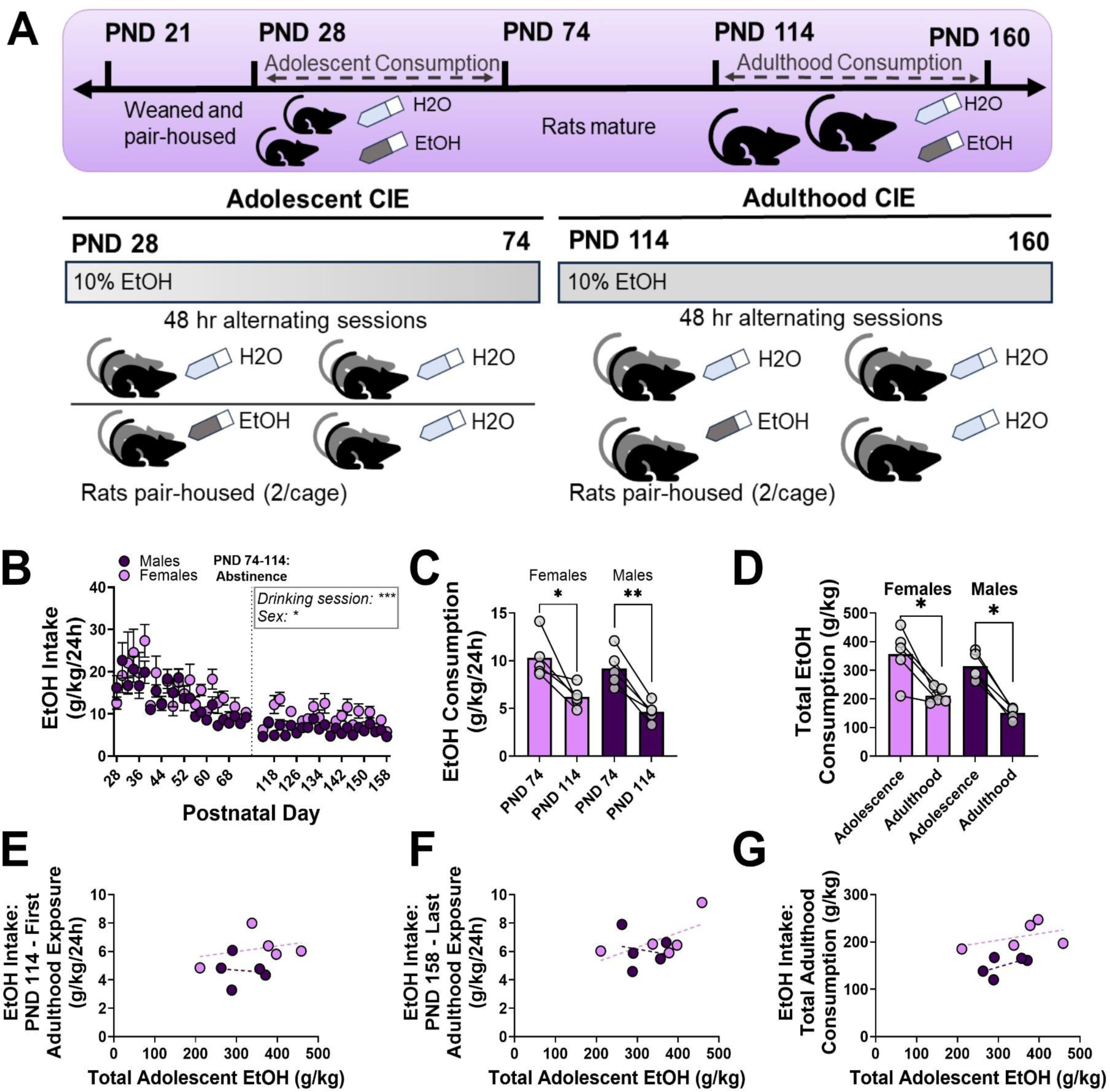
Adolescent CIE does not increase EtOH consumption during adulthood CIE in Fischer 344 rats. (A) Experimental timeline. (B) EtOH intake declines throughout adolescent CIE. (C) Female and male rats show reduced EtOH intake during the first adulthood CIE session on PND 114 compared to the final adolescent CIE session on PND 74. (D) Female and male rats consume less total EtOH during adulthood CIE than in adolescent CIE. (E-G) Neither sex nor total adolescent EtOH intake were significant predictors of adulthood drinking measurements. Trendlines in E-G represent the line of best fit for each sex. * = *p* < .05; ** = *p* < .01; *** = *p* < .001. C-G: Each value represents EtOH intake in one cage of 2 rats.

## 4.0 DISCUSSION

Our current results show that four drinking models, designed for both rats and mice, and across four labs, three institutes, and two species failed to show appreciable increases in adulthood alcohol consumption. This adds to a mixed literature showing both moderate increases (Strong et al., 2010; Lee et al., 2017; Younis et al., 2019; Van Hees et al., 2022) and no effect (Wooden et al., 2023; Gilpin et al., 2012; Gamble & Diaz, 2020; Nentwig et al., 2019) of adolescent alcohol exposure on adulthood drinking in rodent models (for review, see Towner & Varlinskaya, 2020). Importantly, the lack of a robust effect across models suggests that the biological basis of these adolescent drinking effects is nuanced. Our work supports the idea that the positive relationship between adolescent and adult drinking typically seen in humans may be driven by various complex biological and social factors that may not be completely modeled in rodents.

It must be noted that the various rat and mouse models are not identical, thus limiting direct species comparisons across rodents. To this point, it is also possible we are seeing some ceiling effects in experiments 1 and 2, using adolescent DID in mice, due to the confounding stress of social isolation during early adolescence, as single housing stress has been shown to increase alcohol consumption in various rodent models (Lodha, Brocato, & Wolstenholme, 2022; Skelly et al., 2015). Other reports indicate more modest effects of social isolation on EtOH consumption, especially in females (Butler et al., 2014). Additional studies have suggested that adolescent social isolation alone is not enough to increase adulthood alcohol intake, rather the combination of adolescent alcohol exposure with social isolation (Chandler et al., 2022), though we saw only a modest increase in total binge alcohol consumption in single-housed mice exposed to adolescent alcohol compared to single-housed mice which drank only water. Two of the models used (experiment 3, Figure 4; experiment 5, Figure 6) explicitly addressed this concern with a paired housing model. In addition, experiment 4 (Figure 5) utilized mating cages that were divided with perforated clear plastic dividers to minimize isolation stress during 2 hr drinking sessions. Therefore, the effect of isolation is minimized in these three models and likely does not synergize with adolescent ethanol consumption. On this note, limited studies have assessed the combined effects of adolescent alcohol and social isolation exposure; those studies have suggested that social isolation may be the primary driver of any changes in alcohol intake or preference (though some studies show increased preference after social isolation and others show decreased preference) (Schenk et al., 1990; Pisu et al., 2011; for review, see Lodha, Brocato, & Wolstenholme, 2022). Notably, other studies also report increased alcohol consumption in group-housed adolescent mice compared to single-housed mice, particularly in male mice and rats (Logue et al., 2014; Varlinskaya, Truxell, & Spear, 2015). Adolescent isolation stress can interact with other anxiety- and reward-related behaviors (Chappell et al., 2013; Caruso et al., 2018; Walker et al., 2019; Hong et al., 2012) that may interact with adulthood alcohol consumption in differing models. In addition, human adolescent consumption of alcohol often occurs in combination with other substances, including both prescribed (Crowley et al., 2014) and illicit (Torrens et al., 2023) substances that have the potential to interact with alcohol. Importantly, human adolescents are often co-abusing these substances with alcohol (Claus et al., 2022), driving a greater necessity to understand their individual and synergistic effects on brain development. In addition, most of our models used relatively similar adolescent exposure windows that spanned puberty in both sexes across the two species; alcohol consumption or changes in housing conditions during both earlier and later adolescent timepoints may produce different results.

We and others have previously published work showing prefrontal cortical circuitry changes following adolescent binge drinking in rodent models (Sicher et al., 2023; Galaj et al., 2020; Centanni et al., 2017) suggesting that despite the lack of increased drinking seen in the current study, adolescent binge drinking can still produce profound neuroadaptations. In our previous work, electrophysiological differences did not emerge until this longer term timepoint. Immediately after adolescent DID, the effects in males and females were the same (pyramidal neurons were hypoexcitable and SST cells hyperexcitable), with sex differences emerging 30 days after adolescent alcohol exposure (while SST neurons remained hyperexcitable in both sexes, there was a compensatory hyperexcitability of female pyramidal neurons, with a return to control firing of pyramidal neurons in males). Others have found adolescent alcohol consumption interferes with pyramidal neuron membrane properties (Salling et al., 2018). Multiple lines of work have implicated HCN1 channels in the observed prefrontal dysfunction following adolescent alcohol (Salling and Harrison, 2020; Hughes et al., 2020). Together, this suggests that adolescent alcohol consumption is still interfering with prefrontal cortical development. Our study complements work by others showing lasting changes in alcohol sensitivity, cognition, and other AUD-associated phenotypes after adolescent alcohol exposure (Seemiller et al., 2023; Crews et al., 2016; Broadwater & Spear, 2013; Pascual et al., 2007; Wolstenholme et al., 2017), highlighting the nuance in studying these addiction-relevant behaviors.

In the present study, each adolescent alcohol paradigm initiated alcohol consumption at roughly the same age in early adolescence. The current body of work further highlights the need to consider precise developmental timing of alcohol exposure, as adolescence may span multiple periods that are uniquely vulnerable to insults like stress and alcohol use (Spear, 2015; Spear, 2020; Ortelli et al., 2023). Although we did not find robust escalations in alcohol consumption in our preclinical models of voluntary adolescent alcohol exposure, earlier initiation of alcohol use is associated clinically with negative alcohol-related outcomes later in life, including risk of AUD development and other problematic alcohol consumption patterns (Grant & Dawson, 1997; DeWit et al., 2000; Hingson et al., 2006; Patrick et al., 2023). Preclinical research has begun to parse consequences of early versus late adolescent alcohol exposure. For example, intermittent alcohol exposure models in rats have produced unique effects on cognition and alcohol conditioned taste aversion depending on whether exposure occurred in early or late adolescence (Sanchez-Roige et al., 2014; Saalfield & Spear, 2015). Such findings support that there may be separable periods of vulnerability during adolescence (Spear 2015) or throughout the lifespan that may differentially influence later alcohol use patterns. Further, the effects of cumulative alcohol exposure beyond adolescence and throughout the lifespan are important to consider (Seemiller et al., 2024).

Additional social factors can influence patterns of alcohol drinking in adolescents and young adults such as comorbid psychiatric conditions, social norms and policies concerning alcohol use in young people, parental attitudes towards alcohol – which may influence access to alcohol for those below the legal drinking age (Vashishtha et al., 2019; Ning et al., 2020). These factors are difficult to capture in current preclinical models. Preclinical models provide control over experimental conditions, which facilitates the identification of neurobiological consequences of adolescent alcohol use. However, our findings here suggest that these models may not accurately capture early alcohol exposure as a risk factor for higher levels of alcohol consumption later in life.

## 6.0 ACKNOWLEDGEMENTS

Funding: F31AA030455 and T32GM10563 to ARS, T32AA025606 to AL, K99AA030609 to VV, F32AA031396 to LS, R37AA017447 and R01AA029841 to MR, R21AA031101 and SUNY Empire Innovation Funds to FPV, P50AA017823 to FPV, TD, and NAC, R01AA029403 and The Huck Institutes of the Life Sciences Endowment Funds to NAC.

